# Superscan: Supervised Single-Cell Annotation

**DOI:** 10.1101/2021.05.20.445014

**Authors:** Carolyn Shasha, Yuan Tian, Florian Mair, Helen E.R. Miller, Raphael Gottardo

## Abstract

Automated cell type annotation of single-cell RNA-seq data has the potential to significantly improve and streamline single cell data analysis, facilitating comparisons and meta-analyses. However, many of the current state-of-the-art techniques suffer from limitations, such as reliance on a single reference dataset or marker gene set, or excessive run times for large datasets. Acquiring high-quality labeled data to use as a reference can be challenging. With CITE-seq, surface protein expression of cells can be directly measured in addition to the RNA expression, facilitating cell type annotation. Here, we compiled and annotated a collection of 16 publicly available CITE-seq datasets. This data was then used as training data to develop Superscan, a supervised machine learning-based prediction model. Using our 16 reference datasets, we benchmarked Superscan and showed that it performs better in terms of both accuracy and speed when compared to other state-of-the-art cell annotation methods. Superscan is pre-trained on a collection of primarily PBMC immune datasets; however, additional data and cell types can be easily added to the training data for further improvement. Finally, we used Superscan to reanalyze a previously published dataset, demonstrating its applicability even when the dataset includes cell types that are missing from the training set.

## Introduction

The emergence of single-cell genomics in recent years has enabled new levels of precision in analysis of the immune system, providing a deeper understanding of disease on a cellular level and furthering development of immunotherapy treatments and other therapeutics^1–5^. A fundamental component of single cell analysis is classification of the individual cells according to phenotype and developmental stage. This is typically an essential step for subsequent analysis, and has consequently been the focus of significant attention^3,6,7^.

However, differentiation of immune cell types is often challenging to do from gene expression profiling alone (i.e. single-cell RNA sequencing, or scRNA-seq), as functional differences in cells are often not fully reflected in the transcriptome^8,9^. Correlation between gene expression and protein levels is not perfect; the Pearson correlation coefficient between mRNA expression and protein expression has been measured to be between 0.4^10^ and 0.6^11^. A number of computational methods for cell type annotation have been developed in recent years, many of which rely on mapping to a reference genome with pre-labeled cells, for example SingleR^12^, scMatch^13^, and SciBet^14^. However, the significant heterogeneity that can exist between datasets, particularly in the presence of disease, can limit the limit the range of applicability of these methods, and the generation of additional high-quality reference data with labeled cells can be challenging and expensive^3,6^.

Other classification methods rely on a pre-defined set of marker genes for cell type annotation; for example Garnett^15^, CellAssign^16^, and SCSA^17^. While this improves performance in some cases, the reliance on canonical marker genes can be problematic, especially since many of those are based on published data from protein measurements which, as mentioned above, do not correlate directly with gene expression. While curated collections of marker genes for known cell types do exist^18,19^, they are not standardized or universal, and many cell types share marker genes, complicating the classification process. Classification tools that rely on marker genes often require implementation of an unsupervised clustering step first. SCSA, for example, takes the pre-defined clusters as input then uses the top differentially expressed genes in each cluster to annotate the cells based on similarity to a provided list of marker genes. Some reference-based methods include a necessary clustering step first as well; for example CellO^20^, which then annotates clusters using a model trained on reference bulk RNA-seq data. The requirement of an initial clustering step may cause problems in specific cases where datasets are relatively homogenous or when certain cell type populations are very small, in which case the differentially expressed genes may not map well to canonical marker genes.

More recently, a number of multimodal single cell omics approaches have been developed, which allow integration and mapping of multiple data types on the single cell level^21^. These include, for example, T cell receptor (TCR) profiling, spatial transcriptomics, measurement of chromatin accessibility with single-cell ATAC-seq^22–24^, and profiling of protein expression in single cells with Abseq^25^ and CITE-seq^26–28^. CITE-seq (Cellular Indexing of Transcriptomes and Epitopes by sequencing), which uses oligonucleotide-labeled antibodies to measure surface protein expression of single cells, provides additional detail about cell function and phenotype independently of the transcriptome, enabling easier cell annotation based on direct protein expression. mRNA transcript expression is typically much lower, by several orders of magnitude, than protein expression^11,29^, and also has a much smaller range of expression; transcript copy numbers typically only span about two orders of magnitude, while protein copy numbers can span 6-7 orders of magnitude^10,29^. For these reasons, as well as the imperfect correlation between mRNA expression and protein levels, techniques such as Abseq and CITE-seq improve on scRNA-seq for phenotype identification.

While CITE-seq is still a relatively new technique, it is being rapidly adopted by the single cell community. Consequently, more and more datasets containing integrated proteomics data are being published and made publicly available online, although the number and variety of proteins measured in each dataset can vary significantly. When sufficient antibodies are used for CITE-seq measurements, granular cell phenotypes can be determined on a single cell level by gating cells according to cell surface protein expression, as is typically done with flow cytometry. CITE-seq therefore allows easier comparisons of scRNA-seq data to published work that only uses protein markers, as in flow cytometry. Flow cytometry enables direct measurement of surface protein expression through the use of fluorochrome-conjugated antibodies, which can be detected with a light-scattering procedure. Cell populations can then be grouped and identified according to relative protein expression, typically through inspection of two-dimensional plots showing a few proteins at a time. Manual gating is therefore a subjective and often time-consuming process. Efforts have been made to streamline the gating process with the inclusion of automatic techniques, which were reviewed and benchmarked by the FlowCAP Consortium and found to be effective in many cases^30^ when applied to flow cytometry data.

Despite these limitations, the recent availability of public CITE-seq datasets have promising implications for scRNA-seq-based cell classification methods. A recent example of a computational technique that takes advantage of proteomics in cell annotation is Azimuth, a new classification method that uses a reference dataset where labels were generated from integrating proteomics and transcriptomics^31^. Other examples include CITE-sort^32^ and SECANT^33^, both of which take advantage of the antibody-derived tag (ADT) data indicating surface protein expression to annotate cells. However, CITE-seq is still a relatively new technology, and many scRNA-seq datasets do not yet contain protein expression data. Therefore, methods like CITE-sort and SECANT, both of which rely on the inclusion of ADT data to annotate cells, cannot be generalized to datasets containing only RNA information. Azimuth does have this ability, but relies on a certain degree of similarity between the new dataset and the reference in terms of cell type frequency and distribution; it would not, for example, work on a homogenous dataset of a single cell type. This limits its potential for generalization to new tissue or sample types.

As with traditional cytometry data, manual gating on the proteomic component of CITE-seq data can be used to generate reference labels independent of the gene expression data. Our goal is to take advantage of the additional surface marker information from CITE-seq datasets to build a supervised model trained on gene expression data that can annotate RNA-only datasets. In this work, we introduce Superscan (Supervised Single-Cell Annotation): a supervised classification approach built around a simple XGBoost model trained on manually labelled data. Superscan aims to reach high overall performance across a range of datasets by including a large collection of training data. This is in contrast to a method like CaSTLE^34^, which also employs an XGBoost model but requires specification of a sufficiently similar pre-labeled source dataset. To generate the training and testing data, two independent analysts performed manual gating on the corresponding proteomics data from 16 published single cell immune datasets, creating a public resource of several hundred thousand labeled cells that can be used freely for further research.

Superscan is both faster and on average more accurate than the current leading classification methods, as measured against the labels generated manually from ADT data. While the training set currently includes a limited number of cell types, Superscan provides a confidence score along with each prediction that can be used in certain cases to identify cell types that are missing from the training dataset, as we show here with a Merkel cell carcinoma dataset. As more CITE-seq or pre-sorted datasets are generated and become publicly available, they can be added to the training set, expanding the number of included cell types and further improving the model performance.

## Results

### Manual annotation of publicly available CITE-seq datasets

To train an effective supervised classifier, our first step was to assemble as large a collection of datasets as possible that met our criteria: namely, that they were publicly available, included both RNA and ADT raw count matrices, and were of sufficient quality to enable phenotype differentiation based on the ADT assay (see Methods for further details). We collected a total of 16 immune datasets (summarized in Table 1) and manually annotated cells based on the ADT data to generate labels. Manual gating was performed following the framework of Maecker, McCoy, and Nussenblatt 2012. An example of the gating process is shown in Figure 1a. For each cell, broad and fine labels were generated (a list of all labels can be found in Supplementary Table 1), as well as quality scores from 1 (low) to 3 (high) indicating our confidence in each label based on the proteomics data quality.

**Table 1:** Summary of public datasets, including tissue type, number of cells used (i.e. only healthy patients from arunachalam_2020 and su_2020), and number of proteins measured with CITE-seq. Asterisks indicate datasets that were not included in Superscan’s training data.

| name | tissue | # cells | # proteins |
| --- | --- | --- | --- |
| 10X_malt_10k | PBMC | 8412 | 17 |
| 10X_pbmc_10k | PBMC | 8201 | 17 |
| 10X_pbmc_1k | PBMC | 713 | 17 |
| 10X_pbmc_5k_nextgem | PBMC | 5527 | 32 |
| 10X_pbmc_5k_v3 | PBMC | 5247 | 32 |
| arunachalam_2020* | PBMC | 30870 | 39 |
| butler_2019 | BM | 33454 | 25 |
| fournie_2020* | PBMC | 5559 | 12 |
| granja_2019_bmmc | BM | 12602 | 21 |
| granja_2019_pbmc | PBMC | 14804 | 21 |
| hao_2020 | PBMC | 161764 | 224 |
| kotliarov_2020 | PBMC | 58654 | 87 |
| stoeckius_2017 | CBMC | 8617 | 13 |
| su_2020 | PBMC | 44385 | 192 |
| wang_2020 | PBMC | 1372 | 10 |
| witkowski_2020 | BM | 42621 | 84 |

**Figure 1.**
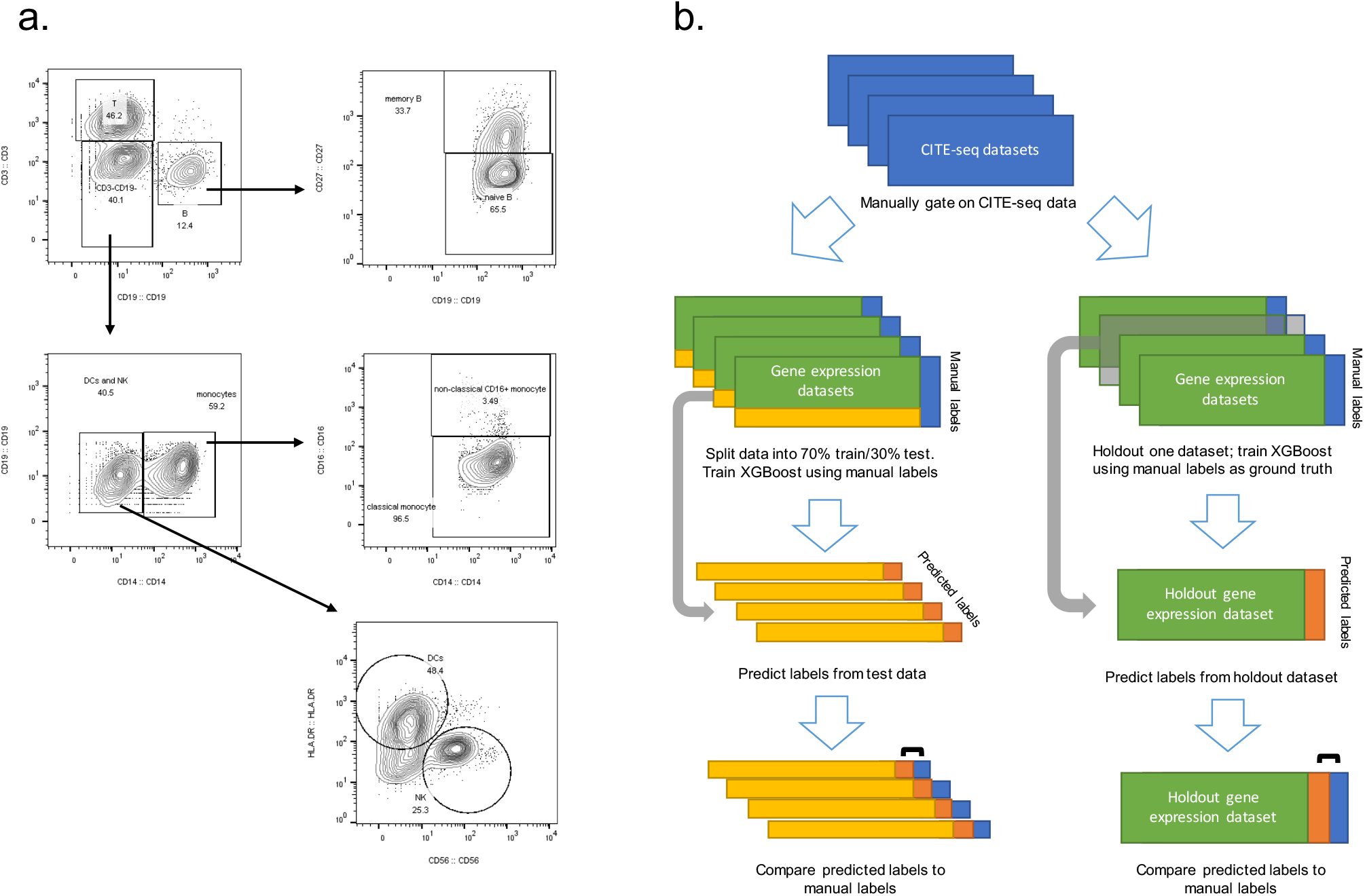
Methodology for manual gating and model training. A. Example bivariate plots showing gating strategy. Gates were drawn in FlowJo according to the protein markers outlined in Supplementary Table 1. B. Methodology for model training, testing, and validation. Data breakdown for initial model development process (feature engineering and parameter optimization) is shown on the left, and the process for validating the model by dataset is shown on the right.

To visually verify the consistency of the manually defined labels, we generated UMAP plots for each dataset. Figure 2 shows several of these example single cell datasets where the UMAPs have been generated based on gene expression (after PCA transform) and surface protein expression alone, colored by the manual labels. Clustering on protein expression data clearly provides superior distinction among immune cell types; CD4 and CD8 T cells, for example, cluster together on the RNA UMAPs because of similar transcriptional profiles, but are separated into distinct groups on the protein UMAPs. This result is expected, given that the manual labels were generated from the protein data, but does reinforce the fact that surface protein expression is able to better distinguish cell phenotypes, motivating our choice to use the ADT data to generate reference labels.

**Figure 2.**
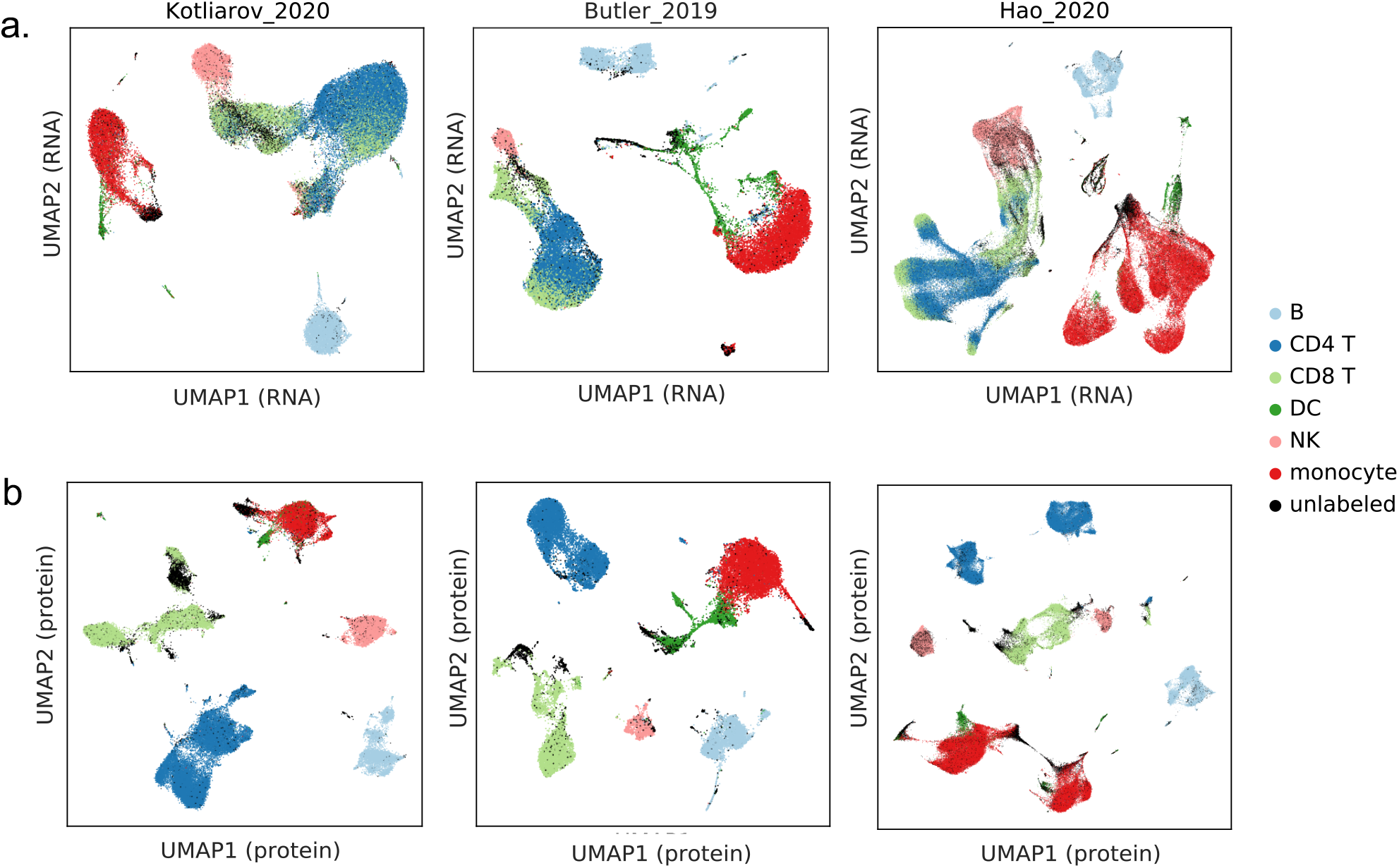
UMAPs of three datasets generated from gene expression and protein expression. UMAPs generated from protein expression result in clearer separation between clusters in all cases. A (top): UMAPs generated from RNA only, colored by manual labels. B (bottom): UMAPs generated from protein data only, colored by manual labels. Three datasets shown (left: kotliarov_2020, middle: butler_2019, right: hao_2020).

### Supervised annotations from RNA can recapitulate broad phenotypes

Our goal in developing Superscan was to create a supervised prediction model that would be broadly applicable to any immune single-cell dataset with RNA measurements, and that could be expanded upon to include additional cell and tissue types in the future. For this reason, we used as many public datasets that we were able to label based on our manual gating strategy and developed the model so as to be able to easily incorporate additional training datasets as they become available.

The methodology for training and evaluation is outlined in Figure 1b. For feature and parameter optimization, 14 datasets were combined and randomly split into a 70% training set and 30% testing set. The remaining two datasets were removed due to low ADT quality (see Methods for further details). The features and parameters identified at this step were then used to test the model on each dataset: for these 14 datasets, each one was held out individually, and the model was trained on the remaining 13 datasets. To test the two datasets that were originally left out of training (arunachalam_2020 and fournie_2020), the model was trained on all 14 original datasets.

Results for broad and fine labels from this holdout procedure are shown in Figure 3a. As expected, overall accuracy when predicting broad labels (median accuracy 0.958) is higher than when predicting fine labels (median accuracy 0.857). This is likely partially due to higher categorical representation (with fewer categories, each class will encompass more cells). Also, some datasets did not have enough proteins measured in order to define certain finer phenotypes (see number of proteins measured for each dataset in Table 1), further reducing the number of cells in each class in the training set. We expect that adding additional high-quality training data where finer phenotypes can be clearly distinguished would improve Superscan’s performance on fine labels.

**Figure 3.**
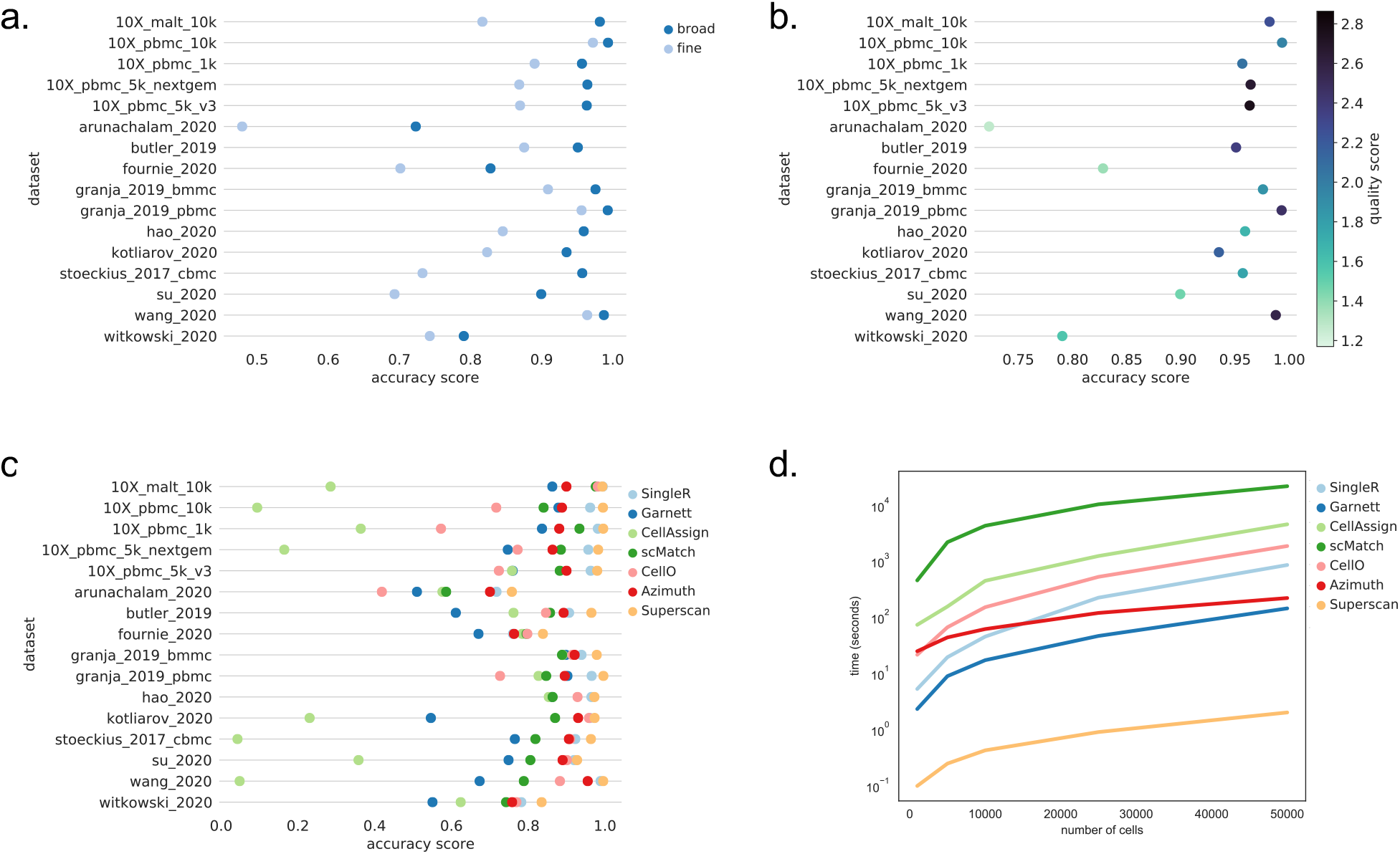
Classification accuracy of Superscan and comparisons to benchmark methods. A. Superscan performance by dataset, broad and fine labels. B. Accuracy of broad labels by dataset, colored by mean quality score of all cells in the dataset. C. Accuracy scores by dataset, colored by cell annotation methods. D. Run time for each cell annotation method, as a function of dataset size (data randomly downsampled from su_2020).

Figure 3b shows the broad label accuracy per dataset, colored by the average quality score (calculated by averaging the quality score for every labeled cell per dataset). As already mentioned, arunachalam_2020 and fournie_2020 have the lowest overall average quality scores, which is why they were removed from the training data. In general, lower quality scores tend to correlate with lower accuracy scores. For many of the datasets, the overall average quality score was lower for the fine labels than for the broad labels (see Supplementary Figure 1a). This may be due to more noise in the data for the necessary markers, e.g. caused by poor antibody staining quality.

### Superscan out-performs competing algorithms in terms of speed and accuracy

Quite a few computational cell annotation packages have been developed in recent years; here, we compared some of the most common and more recent classifiers to our XGBoost-based model, chosen in part based on benchmarking performed in Refs. 7 & 17. The classifiers that we tested were SingleR^12^, Garnett ^5^, CellAssign^16^, CellO^20^, scMatch^13^, and Azimuth^31^. Since each classifier will output different cell labels to varying degrees of specificity, here we compared only broad labels, and mapped classes to general B cells, T cells, dendritic cells (DCs), natural killer (NK) cells, or monocytes. Labels that could not be generalized to these categories were relabeled as ‘other’. Accuracy scores for each classifier and dataset are shown in Figure 3c. Superscan outperformed all other classifiers; SingleR and Azimuth also performed consistently well in general.

To assess run time, one of the larger datasets (su_2020) was downsampled and classifier speed was measured. Results are shown in Figure 3d. Superscan is faster than all others by more than an order of magnitude. In this case, we only compare up to 50,000 cells; on the machine used, datasets containing 100,000 cells or greater had to be broken down into smaller subsets in order to be run when some of the classifiers. This is not an issue for Superscan; when run on a dataset of 1 million cells, classification was completed in just over 2 minutes (122.1s), indicating excellent scalability with dataset size.

Figure 4a illustrates the most important features from the XGBoost model for each broad cell type (see equivalent for fine cell types in Supplementary Figure 2). Reassuringly, many of the top genes correspond to well-known marker genes. Looking at B cells, for example, high gene expression values of MS4A1 (CD20) and CD79A, both known markers for B cells, have a high median SHAP value, indicating that high expression of these genes will push the model towards classifying a given cell as a B cell. Similarly, low values of CD19, another B cell marker, correspond to negative SHAP values, indicating that low CD19 expression discourages B cell classification. Similar clear trends are seen with CD4 and CD8 T cells; low expression of CD8A and CD8B support classification as CD4 T cells, while high expression of CD8A and CD8B support classification as CD8 T cells, as expected.

**Figure 4.**
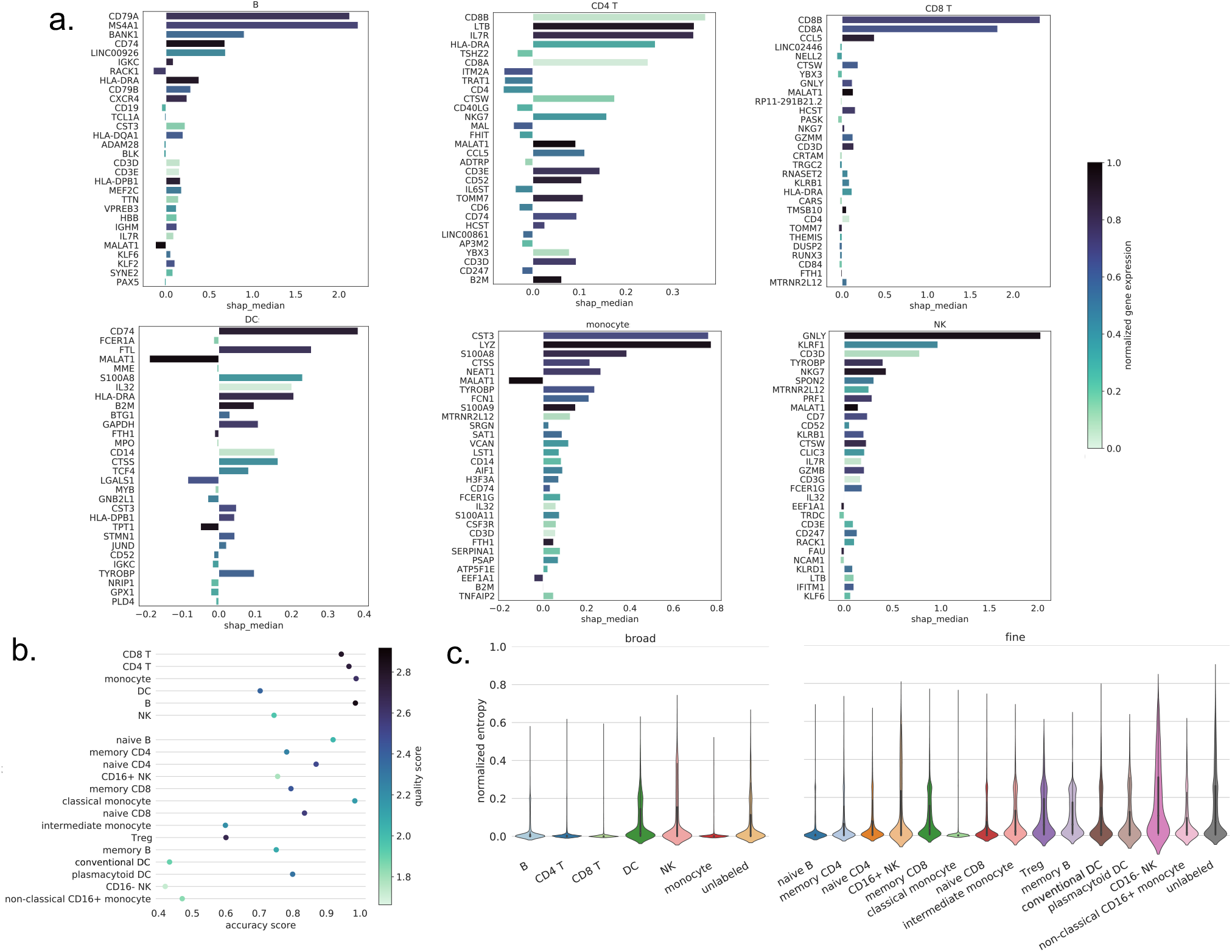
Superscan evaluation metrics by cell type. A. Feature importance for each broad cell type: median SHAP value, ranked by mean(|SHAP value|), colored by standardized expression value log(cpm)/max(log(cpm)). All values averaged over 5-fold CV. B. Accuracy scores (recall) by cell type, colored by mean quality score. C. Normalized entropy scores for each cell type.

### Classification accuracy greatly varies by cell type

We next examine Superscan’s performance by cell type, shown in Table 2, based on a randomized 70/30 train/test split of the data (again removing low quality and unlabeled cells). Looking at the broad labels, dendritic cells clearly have the worst performance, both in terms of precision and recall. However, they also have the lowest representation of all cell types, with fewer than 3,000 DCs out of a total of ∼101,000. B cells have the highest performance, despite comprising only ∼11,000 cells. In general, though, from the fine labels, we can see that cell types that occur less frequently in the data tend to be predicted with lower accuracy. This implies that adding additional high-quality training data containing these less frequent cell types could improve performance in the future.

**Table 2.** Superscan performance by broad (a) and fine (b) cell type, with unlabeled cells removed from training and testing.

| a. | precision | recall | f1-score | support |
| --- | --- | --- | --- | --- |
| B | 0.99 | 0.99 | 0.99 | 10937 |
| CD4 T | 0.97 | 0.97 | 0.97 | 35603 |
| CD8 T | 0.94 | 0.97 | 0.95 | 19960 |
| DC | 0.82 | 0.90 | 0.85 | 2792 |
| NK | 0.96 | 0.94 | 0.95 | 5028 |
| Monocyte | 0.99 | 0.96 | 0.97 | 27517 |
| accuracy |  |  | 0.97 | 101837 |

| **b.** | **precision** | **recall** | **f1-score** | **support** |
| --- | --- | --- | --- | --- |
| CD16+ NK | 0.97 | 0.93 | 0.95 | 4511 |
| CD16- NK | 0.54 | 0.73 | 0.62 | 333 |
| Treg | 0.60 | 0.81 | 0.69 | 1111 |
| Classical monocyte | 0.98 | 0.96 | 0.97 | 25213 |
| Intermediate monocyte | 0.60 | 0.77 | 0.67 | 1101 |
| Memory B | 0.75 | 0.76 | 0.76 | 1655 |
| Memory CD4 | 0.88 | 0.88 | 0.88 | 12948 |
| Memory CD8 | 0.80 | 0.79 | 0.79 | 6660 |
| **Conventional DC** | 0.59 | 0.84 | 0.70 | 851 |
| Naive B | 0.92 | 0.92 | 0.92 | 5830 |
| Naive CD4 | 0.91 | 0.87 | 0.89 | 12652 |
| Naive CD8 | 0.85 | 0.88 | 0.86 | 10253 |
| Non-classical CD16+ monocyte | 0.86 | 0.76 | 0.80 | 827 |
| Plasmacytoid DC | 0.98 | 0.92 | 0.95 | 354 |
| accuracy |  |  | 0.89 | 84299 |

The average accuracy (recall) and quality score by cell type is shown in Figure 4b, now including all cells (not just those with medium or high quality scores). Dendritic cells and NK cells, both of which perform significantly worse than the other broad cell types, have lower average quality scores as well, indicating that labeling of those cell types based on proteomic data quality was more difficult, potentially explaining the worse performance.

In addition to cell classification, the XGBoost model (as implemented in Superscan) can output a vector of probabilities, representing the probability that a given cell belongs to each cell type (these probability values are closely related to the SHAP values shown in Figure 4a). The entropy of these probability values can give an indication of the confidence of the prediction for a given cell. A case where one class has very high probability and the others have low probability would result in a low entropy value, indicating high confidence in the prediction. High entropy values, conversely, indicate lower confidence in a given prediction. Figure 4c shows the distribution of normalized entropy scores (defined in Methods) by cell type. Reassuringly, the lowest performing cell types, e.g. DCs and NK cells, also have the highest average entropy scores, indicating lower confidence in those predictions.

### Superscan enables the annotation of unlabeled cells

So far, we have only looked at classifier performance on cells with defined labels; however, not all cells were able to be manually labeled according to our gating scheme. These unlabeled cells may be cells of entirely different phenotypes, or they may be cells that fall into one of our defined cell types with insufficient or low-quality protein data. Visualizing the predicted cell labels on UMAPs generated from the ADT data (two examples shown in Figure 5a and 5b) can indicate how these unlabeled cells are being classified. Promisingly, the predicted cell labels appear to match well to established clusters. In the kotliarov_2020 data, for example, the unlabeled cells that fall into the monocyte cluster are predicted to be monocytes, and the unlabeled cells in the CD8 T clusters are mostly classified as CD8 T cells. This being said, many unlabeled cells are also higher-entropy, possibly corresponding to cells of different phenotypes (Figure 4c). Looking at the distribution of normalized entropy scores in Figure 5c, cells with higher scores are generally associated with misclassifications, unlabeled cells, and dendritic cells (as seen previously), indicating that the model is less confident in those predictions.

**Figure 5.**
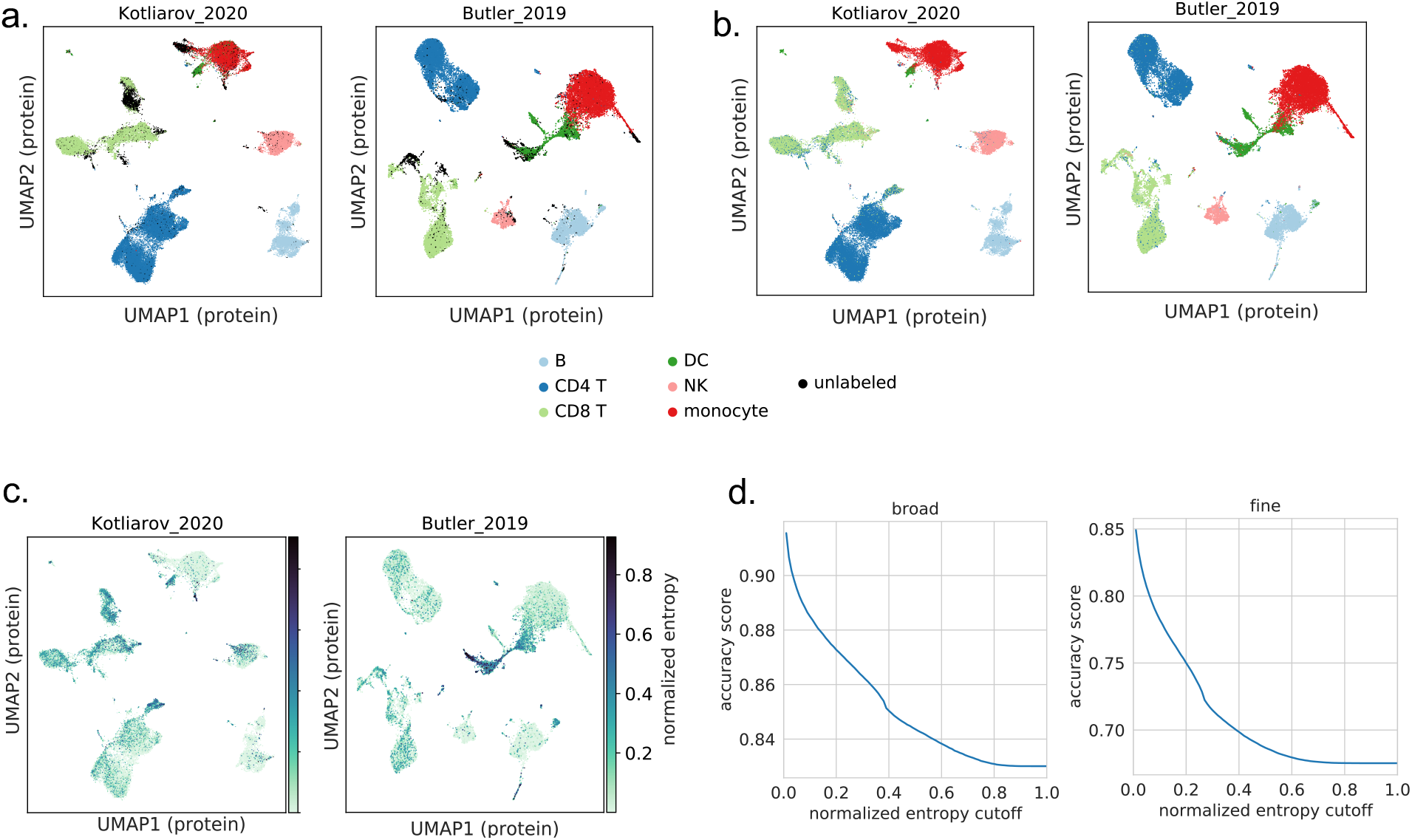
Visualization of Superscan’s performance on unlabeled cells. The normalized entropy score can be used as metric for the model’s confidence in a given prediction. A. UMAPs generated from protein data, colored by manual labels. Two datasets shown (left: kotliarov_2020, right: butler_2019). B. UMAPs generated from protein data, colored by predicted labels from Superscan. C. UMAPs generated from protein data, colored by normalized entropy score from Superscan. D. Accuracy score as a function of normalized entropy cutoff.

### Quantifying prediction uncertainty with entropy scores

To gain more quantitative insight into the connection between the normalized entropy score and accuracy, we consider the effect of incorporating an entropy score cutoff. Figure 5d shows the resulting accuracy score after removing all cells that were predicted with a normalized entropy score above the specified cutoff for a 70/30 train/test split sample, where unlabeled cells are included in the testing set. Unlabeled cells account for 13.3% of the broad labels and 19.5% of the fine labels in the testing set, so the highest possible accuracy when all cells are included (i.e. a normalized entropy cutoff of 1) would be 0.867 and 0.805, respectively. When the normalized entropy cutoff is set to 0.05, the accuracy score rises to ∼0.9 from ∼0.83 and ∼0.8 from ∼0.66 for the broad and fine labels respectively. Reassuringly, this cutoff still preserves 86.6% of the cells (broad labels) and 69.3% of the cells (fine labels). Cells with normalized entropy scores below 0.05 should be considered to be classified with high confidence, while normalized entropy scores between 0.05 and 0.3 indicate moderate confidence, and entropy scores above 0.3 correspond to low confidence. Accordingly, in addition to the normalized entropy score for each prediction, Superscan will also assign a confidence level of “low”, “medium”, or “high” based on the normalized entropy value.

### Entropy scores can be used to identify previously unseen cell-types

We next show an example of how Superscan can be applied to a new scRNA-seq dataset even when it includes cell types that were not in the original training model. We use a Merkel cell carcinoma dataset^35^ with two patients (discovery and validation), which contains a PBMC and tumor sample for each patient (UMAPs of each patient shown in Figure 6a). Superscan predictions are shown in Figures 6b and 6c. Most of the cells from the PBMC samples are classified into fairly well-defined clusters with relatively low entropy scores, with the exception of the cluster of cells indicated with the dashed circle on the UMAPs. These cells are not consistently labeled and have very high normalized entropy scores, indicating low confidence in the predictions. Simple differential expression analysis run on the clusters indicates that these are erythrocytes, a cell type not included in Superscan’s training data.

**Figure 6.**
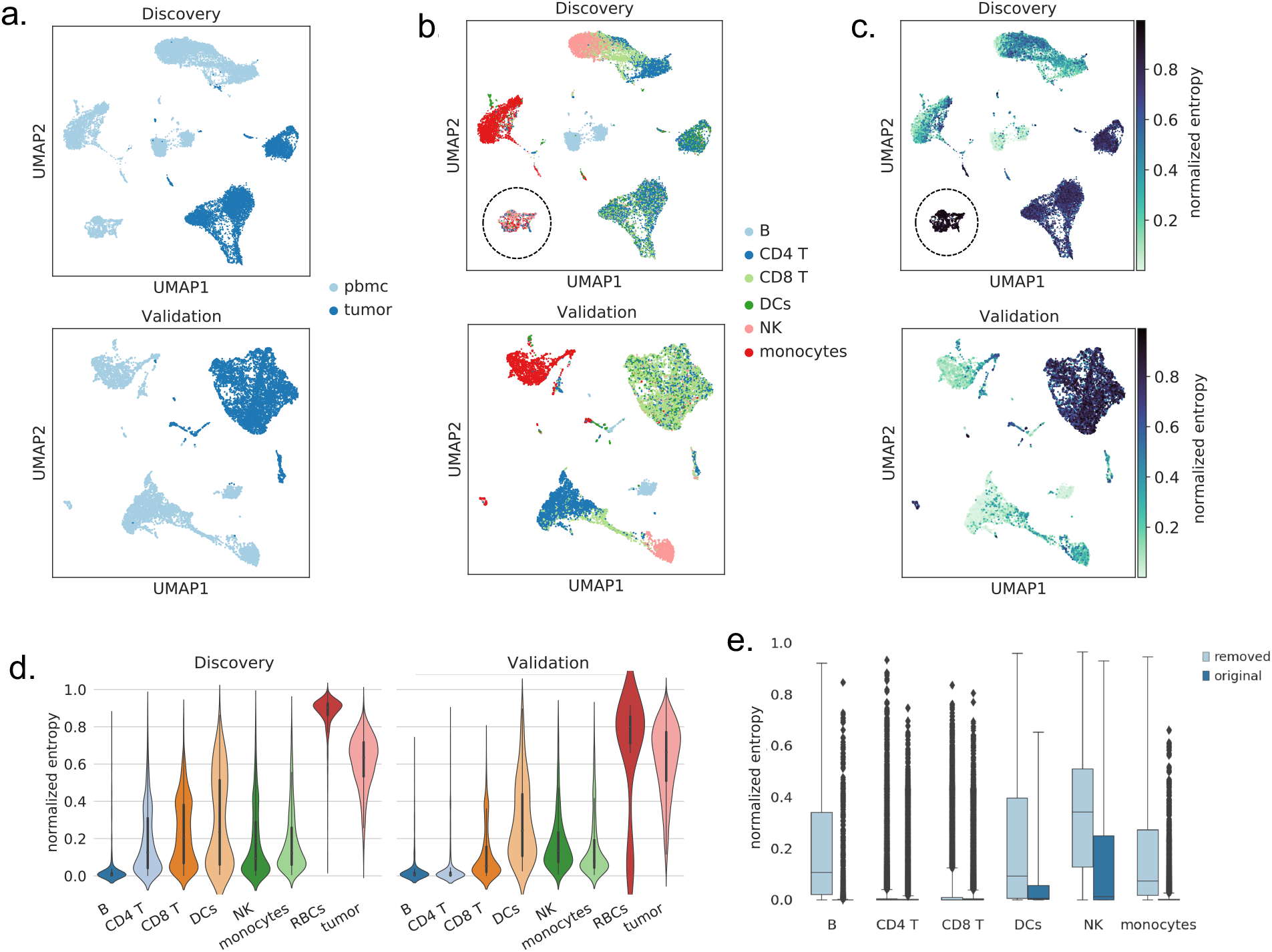
Results from Superscan run on a Merkel cell carcinoma dataset. A. UMAPs of discovery/validation patient, colored by sample type (dataset: paulson_2018). B. UMAPs of discovery/validation patient, colored by Superscan predicted labels. Dashed circle indicates erythrocytes. C. UMAPs of discovery/validation patient, colored by Superscan normalized entropy scores. Dashed circle indicates erythrocytes. D. Normalized entropy scores by cell type for discovery/validation patient. Dashed circle indicates erythrocytes. E. Normalized entropy scores by broad cell type, when cell type was (original) and was not (removed) included in the training data.

Similarly, the large clusters in the tumor samples are also classified with high normalized entropy scores (the other small clusters in the tumor samples are expected to be immune cells in the tumor microenvironment). From this data, then, we can easily identify the clusters that correspond to tumor cells. Labeling these large clusters as tumor cells and the circled cluster as erythrocytes, we can see that the distribution of entropy scores is in fact significantly higher than the other predicted cell types (Figure 6d). Of course, some prior knowledge of the sample composition was required here, but this example demonstrates that in such cases (where one or two cell types are expected in the data but not included in Superscan’s model) the normalized entropy score can be used to identify the missing cell types.

To study further how the model would perform on a new cell type, we can remove one cell type, for example monocytes, from the training data. This would force all monocytes to be classified as one of the other 5 cell types. We can then look at the probability scores for the true monocytes after prediction, shown in Figure 6e. The average entropy is now significantly higher than the other cell types, with very few cells being (mis)labeled with high confidence, with the exception of CD4 and CD8 T cells, which get misclassified (confidently) as each other. A similar situation occurs for fine cells subsets that are part of the same lineage; for example, memory CD8 T cells get classified as mostly naive CD8 T cells, and to a lesser extent memory CD4 T cells, with low uncertainty. However, in most cases, the removed cell type is classified with higher uncertainty (entropy) than in the original model. This confirms that the entropy scores can be used when applying Superscan to datasets that may contain cell types not included here to give some indication of the prediction confidence.

## Discussion

In this work, we took advantage of the recent availability of CITE-seq immune datasets, which provide a means of generating a labeled training set independently of gene expression. The manually labeled datasets are available for public use and exploration. Using this as training data, we developed an automated classifier, Superscan, which is fast, scalable, and easy to use.

An important qualification is that the manual gating performed to obtain the cell labels for training and comparison is necessarily an imperfect process which relies on some degree of subjectivity. To perform manual gating effectively requires a well-defined set of surface markers for each desired phenotype, which can be challenging; the development of a set of universally agreed-upon marker proteins is still ongoing. Even apart from the fact that cell markers for some cell subsets are not well-defined (e.g. regulatory T cells, dendritic cells (DCs), and natural killer (NK) cells), different markers can be used to differentiate well-defined cell subsets, such as the use of CCR7 or CD45RA/CD45RO for defining naive and memory T cells^36^. Finally, deciding whether a cell is positive or negative for a given marker requires choosing a threshold which often is not clearly evident from the data, given the potentially high level of background noise resulting from imperfect antibody staining. Antibody titrating is resource-intensive and therefore commonly not performed, resulting in lower-than-ideal data quality^37^.

Although we attempted to moderate this by including quality scores for each label, it is ultimately impossible to determine a complete ground truth for these datasets. In many cases, datasets do not have sufficient antibodies to determine finer cell subtypes (see Table 1), limiting the comprehensiveness of our labels. The range of performance among datasets seen in Figure 3, therefore, is expected. The average quality scores by dataset for the fine labels, as well as average normalized entropy scores by dataset for both broad and fine labels is shown in Supplementary Figure 1, where it can generally be seen that datasets with lower average quality scores also tend to have lower overall accuracy scores and higher average normalized entropy scores. We do expect that as more high-quality data with higher numbers of markers becomes available, less subjectivity will be required for identifying phenotypes, which should reduce uncertainty in the training set and therefore improve the model.

Superscan is pre-trained on the 14 datasets discussed here and outlined in Table 1; however, we hope to add to the number of training datasets over time. A clear limitation to the current version of Superscan is its inclusion of mostly only PBMC samples and only 6 major immune cell types (due to limited data availability). As shown in Figure 6, inferring new cell types is possible with some pre-existing knowledge, and there is some indication that cells in different sample types (e.g. tumor microenvironment) can also be classified with Superscan. However, we aim to include additional data to the model as high-quality CITE-seq datasets become available, or with pre-sorted datasets. This should ultimately be able to improve the range of applicability, as well as improve overall performance. The poor performance with dendritic cells, for example, is likely at least partially due to the limited number of dendritic cells in the training data. Adding additional high-quality data should therefore improve the performance of Superscan overall.

As Superscan classifies cells on the cell level, it is robust across datasets, regardless of the cell type distribution, for example in a homogenous dataset where all the cells are one type (unlike classifiers such as Azimuth, for example). We also note the scalability of Superscan: it can classify large datasets in a matter of seconds, as the model is pre-trained. It requires 1000 features to run; however, complete overlap with all 1000 features is not necessary since missing values are acceptable within the XGBoost framework. Superscan is freely available for download (see Data Availability).

## Methods

### Dataset preprocessing and manual gating

A total of 16 published CITE-seq datasets, containing both scRNA-seq and ADT data, were collected for training and testing of Superscan. Three datasets were obtained directly from the 10X Genomics “Chromium Demonstration (v3 chemistry)” public collection: 10X_malt_10k (10k Cells from a MALT Tumor - Gene Expression and Cell Surface Protein), 10X_pbmc_10k (10k PBMCs from a Healthy Donor - Gene Expression and Cell Surface Protein), and 10X_pbmc_1k (1k PBMCs from a Healthy Donor - Gene Expression and Cell Surface Protein). Two were obtained from the 10X Genomics “Chromium Next GEM Demonstration (v3.1 Chemistry)” collection: 10X_pbmc_5k_nextgem (5k Peripheral blood mononuclear cells (PBMCs) from a healthy donor with cell surface proteins (Next GEM)) and 10X_pbmc_5k_v3 (5k Peripheral blood mononuclear cells (PBMCs) from a healthy donor with cell surface proteins (v3 chemistry)). The remaining datasets were from the following publications: arunachalam_2020^38^, butler_2019^39^, fournie_2020^40^, granja_2019_bmmc & granja_2019_pbmc^41^, hao_2020^31^, kotliarov_2020^42^, stoeckius_2017^26^, su_2020^43^, wang_2020^44^, and witkowski_2020^45^. No additional preprocessing or normalization was performed.

All datasets were comprised of immune cells, sampled from mostly peripheral blood mononuclear cells (PBMCs), as well as some cord blood mononuclear cells (CBMCs) and bone marrow samples. Details for each dataset, including the number of cells and proteins measured, can be found in Table 1. Two of the datasets (arunachalam_2020 and su_2020) contained data from COVID-19 patients as well as from healthy donors; in those cases, only the data from healthy patients was kept, to avoid severe heterogeneity in the immune system arising from disease. This was also done to avoid over-weighting any one particular dataset (the su_2020 dataset has over 500,000 cells). The number of proteins measured varies significantly among datasets, affecting the granularity of cell phenotypes that could be identified.

Manual gating was performed and verified by two independent analysts following Ref. 36, and as partially shown in Figure 1 (larger version in Supplementary Figure 3). Gating was performed on ArcSinh-transformed ADT raw counts using FlowJo software (FlowJo, LLC). Markers used to gate each cell type are shown in Supplementary Table 1. Two levels of cell labels (broad and fine) were generated for each dataset. Only immune cell types were identified: B cells, CD4 and CD8 T cells, dendritic cells, monocytes, and NK cells. Cells that did not meet the gating criteria for any of these cell types were left as ‘unlabeled’ and removed from the training data. This occurred when the proteomics data was not sufficient to identify the cell type (or when the cell type fell outside of our scope of immune cells). For example, according to our gating scheme (Supplementary Table 1), CD27 is used to differentiate between naive B and memory B cells. However, not all datasets contain ADT data for CD27 (stoeckius_2017, for example). B cells in those datasets therefore will not have a fine label.

For each label, a quality score was assigned that reflected the confidence of our label, based on antibody staining quality, number of cells, and clear separation between populations. The number of antibodies measured (see Table 1) as well as the staining quality varied significantly across datasets, so our confidence in the gating (reflected by the quality scores) varied accordingly. Quality scores ranged from 1 to 3, with 3 corresponding to high quality/confidence. Scores of 3 were given to cells where there were clear bimodal distributions of well-defined subsets (based on protein expression), allowing us to clearly delineate a well-defined biological population. A score of 1 was assigned when the protein distribution was not bimodal, making the gate difficult to determine, or when a gate encompassed less than 10 cells. Scores of 1 were also assigned when the staining was of poor quality and therefore expression levels overall were low. Quality scores of 2 were assigned for cases that fell in between the criteria set for 1 and 3.

### XGBoost implementation

XGBoost (eXtreme Gradient Boosting) is an ensemble, regression-tree based model developed several years ago^46^. Here, we use the Python implementation of XGBoost, with the scikit-learn wrapper^47^. The model was trained on a data frame constructed from the raw gene expression counts matrix; no preprocessing was required. Parameter optimization was performed with the scikit-learn randomized search function, using 3-fold cross-validation; final parameters used were gamma = 0, max_depth = 11, reg_lambda = 2, eta = 0.4, alpha = 0, and n_estimators = 350. The top 1000 features were extracted using the scikit-learn *feature_importances_* attribute. SHAP (SHapley Additive Explanations) values were computed using the shap Python package^48^. Probability vectors for each prediction were extracted using the scikit-learn *predict_proba* function, and entropy scores of each vector were calculated. By normalizing the entropy by log(n), where n is the total number of classes included in the model, we have a standardized value, the normalized entropy, (also called efficiency), which can be used to compare prediction confidence across groups.

### Initial model development and optimization

For initial training and testing of the model, we removed two of the 16 datasets, arunachalam_2020 and fournie_2020, which had the lowest overall proteomics data quality (based on our assignment of quality scores). We also removed all cells with a quality score of 1 (low), and cells taken from COVID-19 patients (as mentioned previously). This left approximately 340,000 cells. For initial development and optimization of the model, we took a random subset of 70% of the cells to use as training, with the remaining 30% held out for testing. Five-fold cross validation was performed. Seven pre-built classification models were initially tested: XGBoost, Random Forest, Decision Tree, Extra Tree, AdaBoost, SVC (support vector classifier), k-nearest neighbors (KNN), and a Multilayer Perceptron (MLP) neural net (see Supplementary Figure 4). XGBoost was chosen as the final model due to its superior performance on both accuracy and run time.

Feature selection was done by selecting a subset of genes based on the feature importance output from the XGBoost classifier and removing mitochondrial and ribosomal genes. We found that using the top 1000 features gave the best performance (in addition to significantly speeding up the classifier). We then performed a randomized search to determine the optimal parameters. Both of the above steps were performed using the 70/30 train/test split of the data as described above, and with 5-fold cross-validation. This was done separately for the broad labels and the fine labels, resulting in an optimized set of parameters for the untrained (at this point) model.

To test performance by dataset, for the original 14 datasets, the model was then trained on the remaining 13 datasets, with the held-out dataset used for validation. To test the remaining two datasets (arunachalam_2020 and fournie_2020), the model was trained on all 14 original datasets, which produced the final version of Superscan. Cells with low quality scores were removed from the training data in all cases, as described above, but testing was performed on all labeled cells. For validation, the model was evaluated on overall classification accuracy: the percentage of cells labeled correctly. For each class, precision and recall were reported in order to examine the model’s performance when predicting less frequent cell types. Consistent performance across different datasets is one of our evaluation criteria as well.

Since Superscan is pre-trained, classification of new data can be done extraordinarily quickly. Superscan requires 1000 specific genes for classification which can be easily subset from any new dataset, meaning training of a new model based on available genes isn’t necessary, unlike with classifiers such as CellO. Furthermore, since XGBoost can easily handle missing values, it is not strictly necessary that a new dataset to be annotated contain all 1000 genes.

### Comparison to other cell type labeling tools

For each cell annotation tool tested, any required data preprocessing as outlined by the relevant package was performed (e.g log-normalization, computation of size factors). Default immune reference datasets or marker gene sets were chosen where necessary: the Monaco immune dataset reference was used for SingleR^49^, and the default Functional Annotation Of The Mammalian Genome 5 (FANTOM5) reference^50^ was used for scMatch. For Garnett, the pre-trained PBMC classifier and marker file were used (hsPBMC), and the default marker genes were used for CellAssign. The hao_2020 dataset is the reference data used for Azimuth, and so that point is excluded. All classifiers were run on the single-cell level, where possible, and default parameters were used (where applicable). In measuring computational speed, only the actual classification step or function was measured, where possible. All classifiers were run on an 8-core machine. In the case of CellO, the model was pre-trained, so that the measured execution time did not include model training.

## Data availability

Superscan was implemented in Python3, and is available on Github: https://github.com/cshasha/superscan. Labeled datasets can be downloaded from https://fh-pi-gottardo-r-eco-public.s3.amazonaws.com/SingleCellDatasets/SingleCellDatasets.html.

## Acknowledgements

We acknowledge the Scientific Computing Infrastructure at Fred Hutch funded by ORIP grant S10OD028685, the J. Orin Edson Foundation, the Translational Data Science Integrated Research Center of the Fred Hutch, and NIH U19AI128914.

## Author Contributions

C.S. developed the code for Superscan, performed the analysis, and wrote the manuscript. Y.T. collected the public datasets, and Y.T. and F.M. performed manual gating of the datasets. H.M. provided input and assisted with testing Superscan. R.G. was the principal investigator, provided direction throughout the project and helped write the manuscript.

