## Supplement for "Superscan: Supervised Single-Cell Annotation"

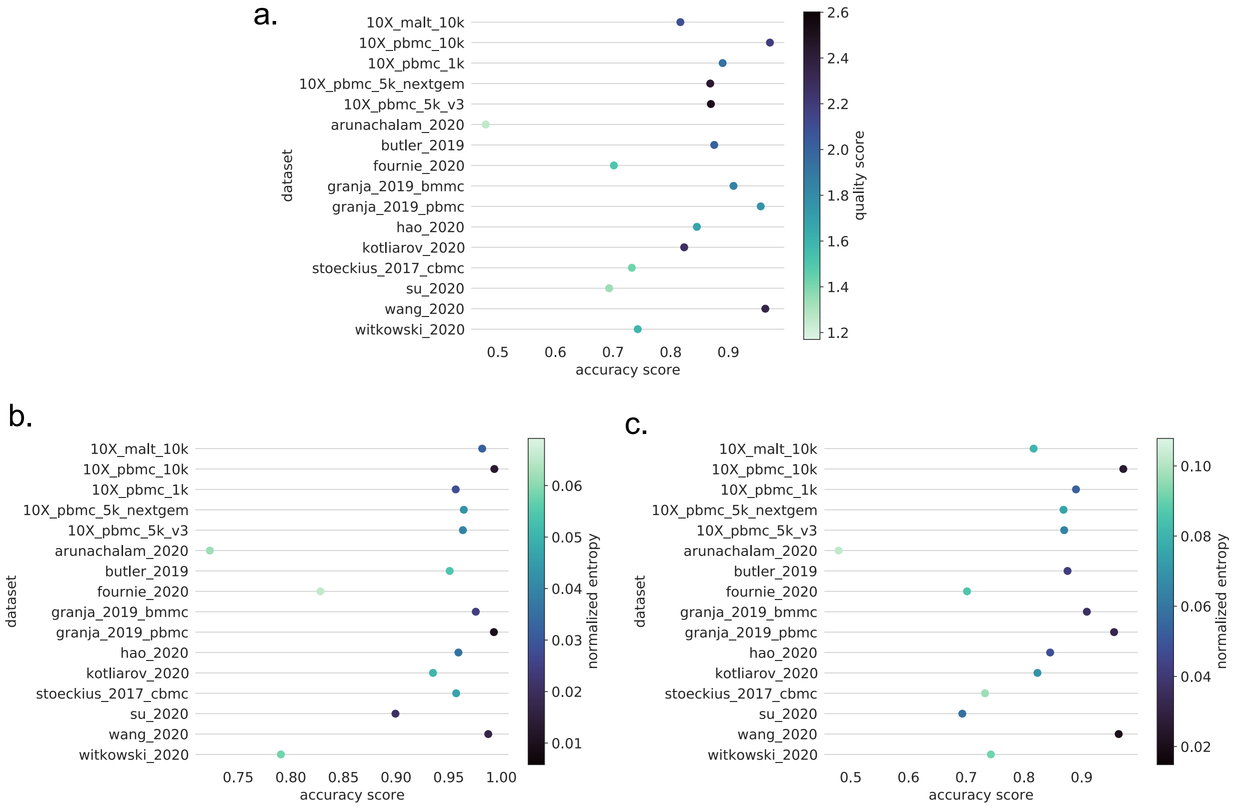


<u>Supplementary Figure 1.</u> A. Accuracy of fine labels by dataset, colored by mean quality score of all cells in the dataset. B. Accuracy of broad labels by dataset, colored by mean normalized entropy of all cells in the dataset. C. Accuracy of fine labels by dataset, colored by mean normalized entropy of all cells in the dataset.

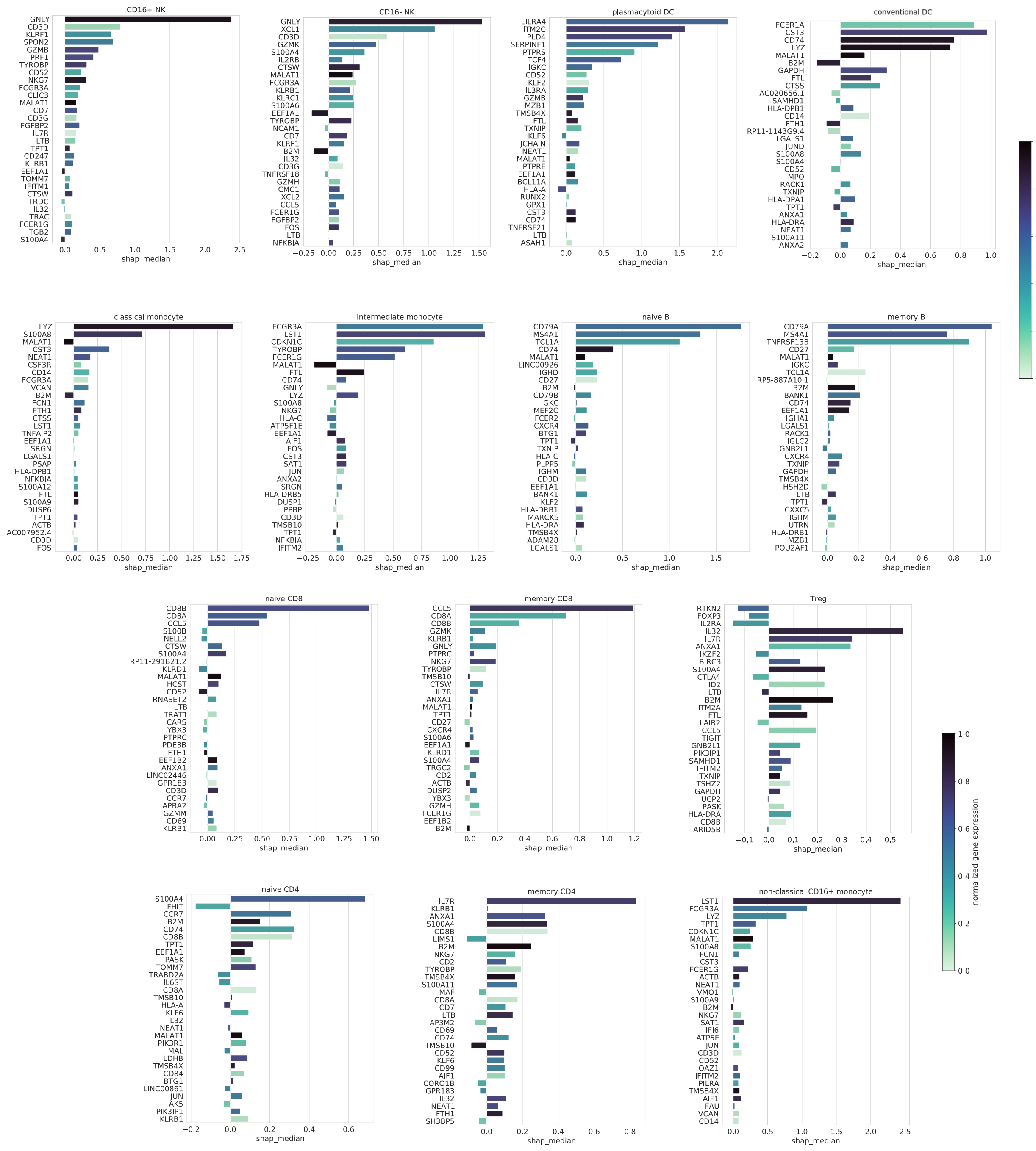


<u>Supplementary Figure 2.</u> Feature importance for each fine cell type: median SHAP value, ranked by mean(|SHAP value|), colored by standardized expression value log(cpm)/max(log(cpm)). All values averaged over 5-fold CV.

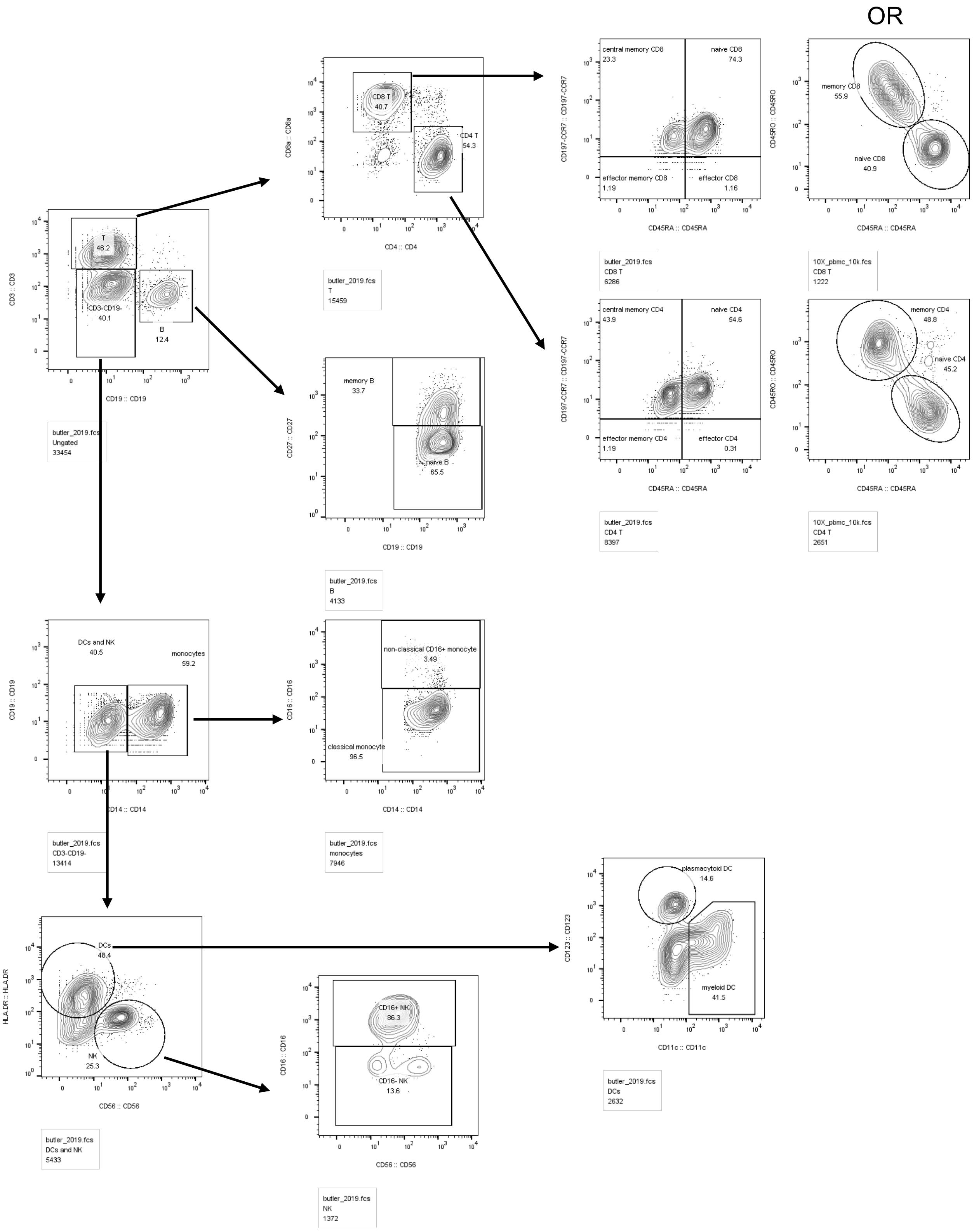


Supplementary Figure 3. Example FlowJo plots showing gating strategy (larger version). Gates were drawn according to the protein markers outlined in Table S1.

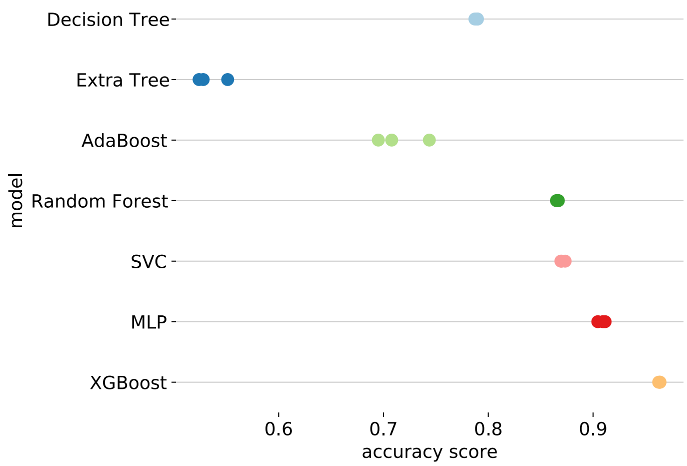


Supplementary Figure 4. Model performances on broad labels with 3-fold cross-validation. For the best-performing models (XGBoost, MLP, and Random Forest), feature and parameter optimization was performed. Scores here reflect the best-performing model; for XGBoost and Random Forest, that included only the top 1000 features. For MLP, several different combinations of layers/dimensions were tested; the best-performing (shown here) used three hidden layers (dimensions 100x50x100), with a ReLU activation function and Adam optimization algorithm. Although SVC had high relative accuracy, the run time was too long to realistically perform optimization. K-nearest neighbors (KNN), not shown, also had a run time that was too long to realistically include.

Supplementary Table 1. Protein markers used for gating each cell type.

| **Cell Type** | **Markers** |
|---|---|
| B | CD3-CD19+ |
| CD4 T | CD3+CD19-CD4+CD8- |
| CD8 T | CD3+CD19-CD4-CD8+ |
| DC | CD3-CD19-CD20-CD14-HLA-DR+CD56-CD16- |
| NK | CD3-CD19-CD20-CD14-HLA-DR-CD56+ |
| Monocyte | CD3-CD19-CD20-CD14+ |
| Naive B | CD3-CD19+CD27- |
| Memory B | CD3-CD19+CD27+ |
| Naive CD4 | CD3+CD19-CD4+CD8-CD25-CD45RA+CD45RO- |
| Memory CD4 | CD3+CD19-CD4+CD8-CD25-CD45RA-CD45RO+ |
| Treg | CD3+CD19-CD4+CD8-CD25+CD127- |
| Naive CD8 | CD3+CD19-CD4-CD8+CD45RA+CD45RO- |
| Memory CD8 | CD3+CD19-CD4-CD8+CD45RA-CD45RO+ |
| Plasmacytoid DC | CD3-CD19-CD20-CD14-HLA-DR+CD56-CD16-CD123+CD11c- |
| Conventional DC | CD3-CD19-CD20-CD14-HLA-DR+CD56-CD16-CD123-CD11c+ |
| CD16+ NK | CD3-CD19-CD20-CD14-HLA-DR-CD56+CD16+ |
| CD16- NK | CD3-CD19-CD20-CD14-HLA-DR-CD56++CD16- |
| Classical monocyte | CD3-CD19-CD20-CD14+CD16- |
| Intermediate monocyte | CD3-CD19-CD20-CD14+CD16+ |
| Non-classical CD16+ monocyte | CD3-CD19-CD20-CD14-HLA-DR+CD56-CD16+ |
